# Topical cream with live lactobacilli modulates the skin microbiome and reduce acne symptoms

**DOI:** 10.1101/463307

**Authors:** Sarah Lebeer, Eline Oerlemans, Ingmar Claes, Sander Wuyts, Tim Henkens, Irina Spacova, Marianne van den Broek, Ines Tuyaerts, Stijn Wittouck, Ilke De Boeck, Camille N. Allonsius, Filip Kiekens, Julien Lambert

**Affiliations:** University of Antwerp, Department of Bioscience Engineering, Groenenborgerlaan 171, B-2020 Antwerp, Belgium; University of Antwerp, Department of Pharmaceutical, Biomedical and Veterinary Sciences, Laboratory of Pharmaceutical Technology and Biopharmacy, Universiteitsplein 1, B-2610 Wilrijk, Belgium; University Hospital Antwerp/University of Antwerp Department of Dermatology and Venereology, Wilrijkstraat 10, 2650 Edegem, Belgium

**Author notes:** Equal contribution (Sarah Lebeer, Eline Oerlemans, Ingmar Claes). **Correspondence:** Sarah Lebeer, University of Antwerp, Department of Bioscience Engineering, Groenenborgerlaan 171, B-2020 Antwerp, Belgium.

## Abstract

The skin is home to an important part of our commensal microbiota, despite it being a cool, acidic and desiccated environment. Tailored microbiome modulation approaches with, for example probiotics, are highly challenging for this body site. Here we show by next-generating sequencing that *Lactobacillus* taxa -especially those known to be dominant in the human vagina- are underestimated members of the skin microbiota. Specific *Lactobacillus* strains were selected in the lab and formulated in a viable form in an oil in water-based topical cream. Facial application by patients with mild-to-moderate acne symptoms was able to reduce inflammatory lesions and comedone formation. This was associated with a temporary modulation of the skin microbiome, including a reduction in relative abundance of staphylococci and an increase in lactobacilli. Skin microbiome modulation by addition of carefully formulated lactobacilli seems to be new therapeutic option to reduce antibiotic use for common acne symptoms.

## Introduction

Being the most extensive interface of the human body with the environment, the skin acts as a home to an important part of our commensal microbiota. Similar to the gut, the skin microbiota have essential roles in the education of our immune system and the protection against invading pathogens and other foreign substances. With recent advances in DNA sequencing approaches, our knowledge has been improved on the biogeography of the skin microbiota at different body sites^1^. We are now transitioning from these descriptive, observational studies towards a better understanding of the functional roles of the commensal microbiota, allowing the design of tailored modulation approaches. However, compared with the richer environment of our intestines, the skin lacks many nutrients beyond basic proteins and lipids, with sweat, sebum and the stratum corneum being main resources^2^. In addition, the skin is a cool, acidic and desiccated environment and skin cells are frequently renewed and shed, so that strategies targeting the skin microbiome are highly challenging. For example, probiotics, i.e. live micro-organisms that, when applied in adequate amounts, promote a health effect on the host^3^, have not yet been widely considered for direct application on the skin.

One of the most common skin diseases is acne vulgaris, a chronic inflammatory skin condition of the sebaceous follicles and glands. The pathogenesis of acne vulgaris is multifactorial, with increased sebum production, alteration in the quality of sebum lipids, dysregulation of the hormone environment and follicular hyperkeratinization as contributing factors. In addition, specific strains of the facultative anaerobe *Cutibacterium acnes* (formerly known as *Propionibacterium acnes*^4^) are involved in the inflammation of the skin, especially by secreting lipase enzymes that are able to metabolize sebum into free fatty acids which may lead to skin irritation^5^. Yet, the observation that almost all adults are colonized with *C. acnes* but only a minority have acne, highlights that other bacteria such as *Staphylococcus* species can be linked to acne pathogenesis as pathobionts or disease modulators^6^. Therefore, both oral and topical antibiotics such as doxycycline, minocycline and clindamycin are frequently used by acne patients^7^, but because of rising problems of antibiotic resistance, various alternative therapies need to be developed^8^.

Here we explored the potential of topically applied, live probiotic lactobacilli to beneficially modulate cutaneous microbial interactions and host inflammatory responses in subjects with mild-to-moderate acne symptoms. Lactobacilli were selected based on their long history of safe use in fermented foods^9^, the gastro-intestinal^10^, urogenital tract^11^ and nasal cavity^12^, but it was unsure whether lactobacilli could also thrive and have health-promoting activities on the skin.

## Results and discussion

### Prevalence of *Lactobacillus* on the skin

Because lactobacilli are not considered to be commensals of the skin, we first monitored the prevalence of lactobacilli on the skin of healthy volunteers. Their relative abundance was explored through 16S amplicon sequencing via Illumina MiSeq (separate runs for V1V2 and V4 variable regions) of facial skin samples (cheek) of 30 volunteers (15 male and 15 female), who did not display acne-related symptoms. In the samples of all female volunteers and 12 male volunteers *Lactobacillus* sequences were found (**Figure 1a)**. *Lactobacillus* species generally did not occur in the top five of most abundant taxa present on the skin. However, some volunteers showed a relative high abundance of *Lactobacillus* taxa (amplicon sequence variants or ASVs), up to 6.4% (based on V1V2 16S sequencing) or 14.3% (by V4 16S sequencing) (**Figure 1b, Extended data Figure 1a-b)**. The relative abundance of *Lactobacillus taxa* based on both runs (V1V2 and V4) was also 10-fold higher in women compared to men, with an average relative abundance of 0.8% (1.4% in women and 0.2% in men, Kruskal-Wallis p= 0.0005). **(Extended Data Figure 1b)**. *Lactobacillus* taxa can thus be considered as endogenous members of the skin microbiota, although their relative abundance is lower than *Staphylococcus, Corynebacterium, Cutibacterium* (often still classified as *Propionibacterium*), and *Streptococcus*, which were the most the dominant taxa in our dataset for both variable regions sequenced **(Extended Data figure 1 a)**.

**Figure 1.**
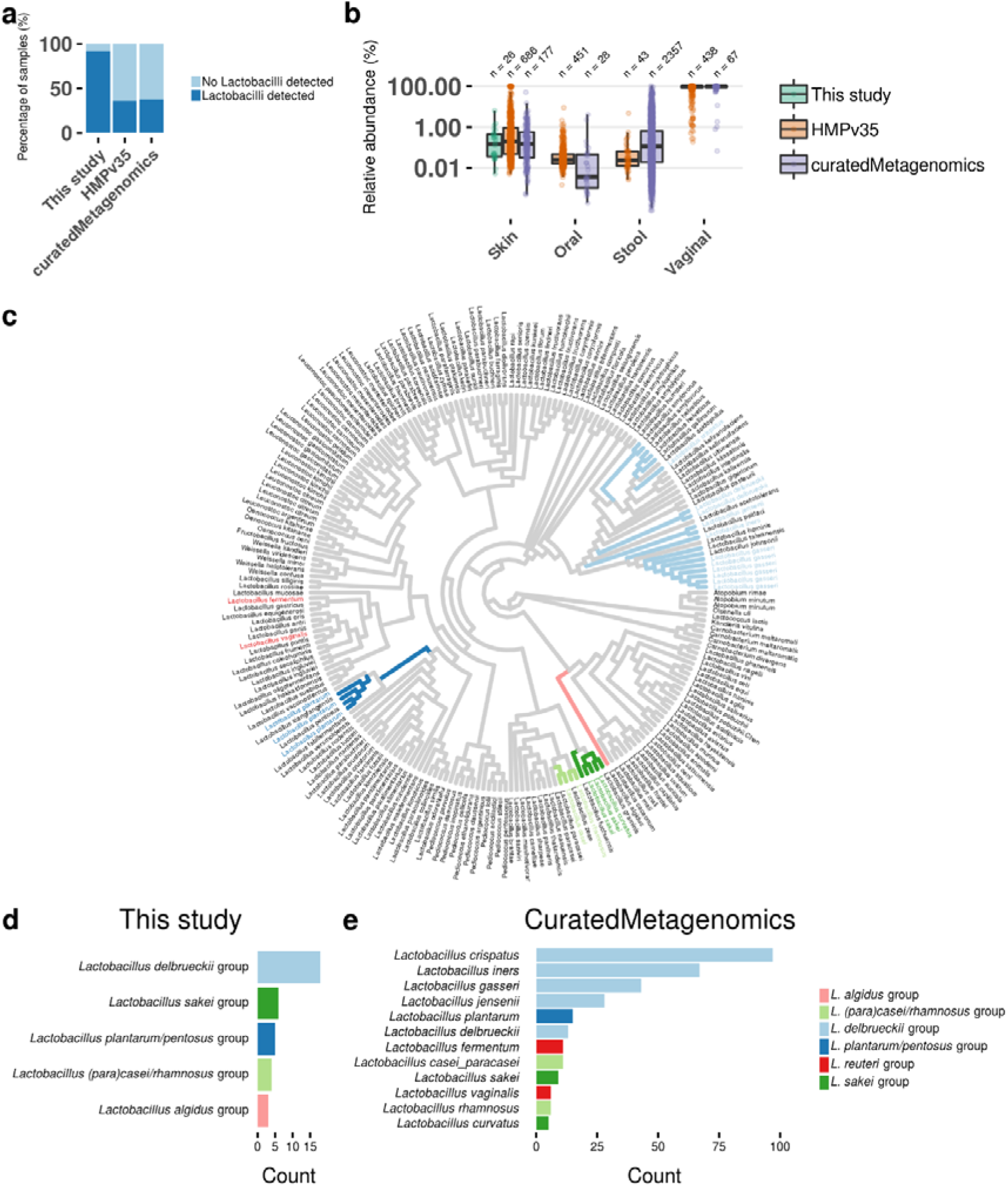
*Lactobacillus* taxa in skin samples of 16S rRNA amplicon and shotgun metagenomic data. (a) Presence and absence of *Lactobacillus* in skin samples of this study, the Human Microbiome Project (HMPv35)^46^ and shotgun metagenomic datasets from three studies^14, 15 and 16^ accessed through the curatedMetaganomics R Package, (b) Comparison of relative abundance of *Lactobacillus* in different niches of the three used datasets. The y-axis is represented in log scale, (c) 16S rRNA cladogram of the *Lactobacillus* Genus Complex. Branches are colored based on phylogenetic placement of Lactobacillus ASVs from this study and the phylogenetic group (as described by Duar *et al.*^19^) they belong to. Tip labels are colored based on the 12 most abundant *Lactobacillus* species found in the skin shotgun metagenomic datasets, (d and e) The most abundant *Lactobacillus* members in this study (d) and the skin shotgun metagenomic datasets (e) colored according to the phylogenetic group of the *Lactobacillus* Genus Complex they belong to.

To confirm our in-house generated data and investigate whether our results are facial site-specific, the presence of lactobacilli was also substantiated in publicly available skin metagenome shotgun datasets by using the curatedMetagenomicData R-package recently described by Pasolli *et al.*^13^. In total, 466 samples from three different studies^14, 15 and 16^ were analyzed. Of these samples, 38% (177/466) showed the presence of at least one *Lactobacillus* species **(Figure 1a)**, but only 29 samples showed a relative abundance higher than 1%. Yet, high relative abundances up to 52% on the skin were also observed (average relative abundance based on curated metagenomics was 3.79%) (**Figure 1b)**. We also included 16S amplicon data from the Human Microbiome Project (V3V5)^17^ where the relative abundance was 24.9% on average due to some outliers having up to 90% relative abundance (**Figure 1b)**. The relative abundance of *Lactobacillus* sequences in the skin samples was also compared to the publicly available data of other human body sites (both 16S and curated metagenome) (**Figure 1b)**. As expected, the vagina showed the highest relative abundance of *Lactobacillus* taxa, but the skin turned out to be the second most important niche for these taxa. Moreover, to have a better idea of the phylogenetic diversity of all *Lactobacillus* taxa present, we also plotted all data on a phylogenetic tree of the *Lactobacillus* genus complex^18^ (**Figure 1c)**. These data indicate that taxa typically associated with the human vagina, *Lactobacillus crispatus, L. iners, L. gasseri* and *L. jensenii* were also found as the most prevalent lactobacilli on the skin (**Figure 1d and 1e**). Also members of the more niche-flexible *Lactobacillus* taxa^19^, *i.e.* from the *L. plantarum/L. pentosus* group and *L. casei/paracasei/rhamnosus* group, were frequently detected **(Figure 1d and 1e)**. The occurrence of *Lactobacillus* taxa on the skin is in agreement with the fact that after normal delivery through the birth canal, these bacteria originating from the mother are among the first to colonize neonate skin^20^. The data presented here (**Figure 1 a-e)** indicate that these *Lactobacillus* taxa are still present in adults, but do not stay dominant in the different human body skin sites studied. Yet, despite their low relative abundance, they could still play a role as keystone microbes, recently redefined as taxa exerting a considerable influence on microbiome structure and functioning irrespective of their abundance across space and time^21^. Therefore, we subsequently aimed to manipulate biotic interactions of lactobacilli on the skin.

### Rationale *in vitro* strain selection

*Lactobacillus* strains were selected from our in-house available laboratory collection (**Extended Data Table 1)** for tailored application in patients with mild-to moderate acne symptoms. A thorough screening approach was applied based on the rationalization that the strains had to be safe, applicable (being robust and showing niche-flexibility as described for lactobacilli by Duar *et al.*^19^) and have the capacity to exert the desired beneficial functions on the human skin including microbiome modulation, immune modulation and epithelial barrier enhancement (**Figure 2a)**. Key properties were substantiated with laboratory tests, genome screening and information available in the literature. Three *Lactobacillus* strains were selected i.e. *Lactobacillus rhamnosus* GG, *Lactobacillus plantarum* WCFS1 and *L. pentosus* KCA1. The rationale for these strains was based on their genome availability^22-24^, knowledge of their host interaction capacity ^25,26^, their robustness and growth capacity^22,24,26^ (**Extended Data Figure 2a)**, in addition to information on their safety in humans after oral^27,28,29^, nasal^30^ and vaginal^31^ high-dose application. *L. rhamnosus* GG was also selected because of previous reports on its capacity to inhibit the toxic effects of *S. aureus* on epidermal keratinocytes^32^, its strain-dependent capacity to promote re-epithelialization^33^ and to augment tight-junction barrier function in human primary epidermal keratinocytes^34^, and our previous experience with this probiotic strain^25^. For microbiome modulation, *C. acnes* was targeted as model pathobiont associated with the inflammatory character of acne vulgaris. *S. aureus* was also targeted as an important pathogen causing skin inflammation. When the activity of spent culture supernatant of our collection of *Lactobacillus* strains was screened for antimicrobial effects on the growth of *C. acnes* in suspension, all *Lactobacillus* strains tested inhibited the growth of *C. acnes* ATCC6919 and *S. aureus* ATCC29213, but *L. pentosus* KCA1 (vaginal origin) and *L. plantarum* WCFS1 (saliva origin) were among the bacteria tested able to exert the highest inhibition **(Figure 2b and Extended Data Figure 2b)**. Other related strains tested, including *Staphylococcus epidermidis* 12228, did not inhibit *C. acnes* growth **(Extended data Figure 2)**. In addition, these strains were able to significantly reduce the lipase activity of *C. acnes* (Figure 2c). These lipase enzymes are involved in inflammation of the skin induced by *C. acnes*, because they metabolize sebum into free fatty acids which may lead to skin irritation^5^. Furthermore, because lactic acid has a strong antimicrobial activity^35^, as well as a documented dose-dependent capacity to ameliorate the appearance of keratoses and acne in dermatology^36^, we also substantiated lactic acid production by the selected lactobacilli (Figure 2d). Furthermore, we validated that the three selected lactobacilli did not exhibit toxic or overt inflammatory responses on primary skin cells (Figure 2e), in agreement with genome predictions^22-24^ and laboratory validation of antibiotic resistance profiles according to the guidelines of the European Food Safety Authority (EFSA)^37^.

**Figure 2.**
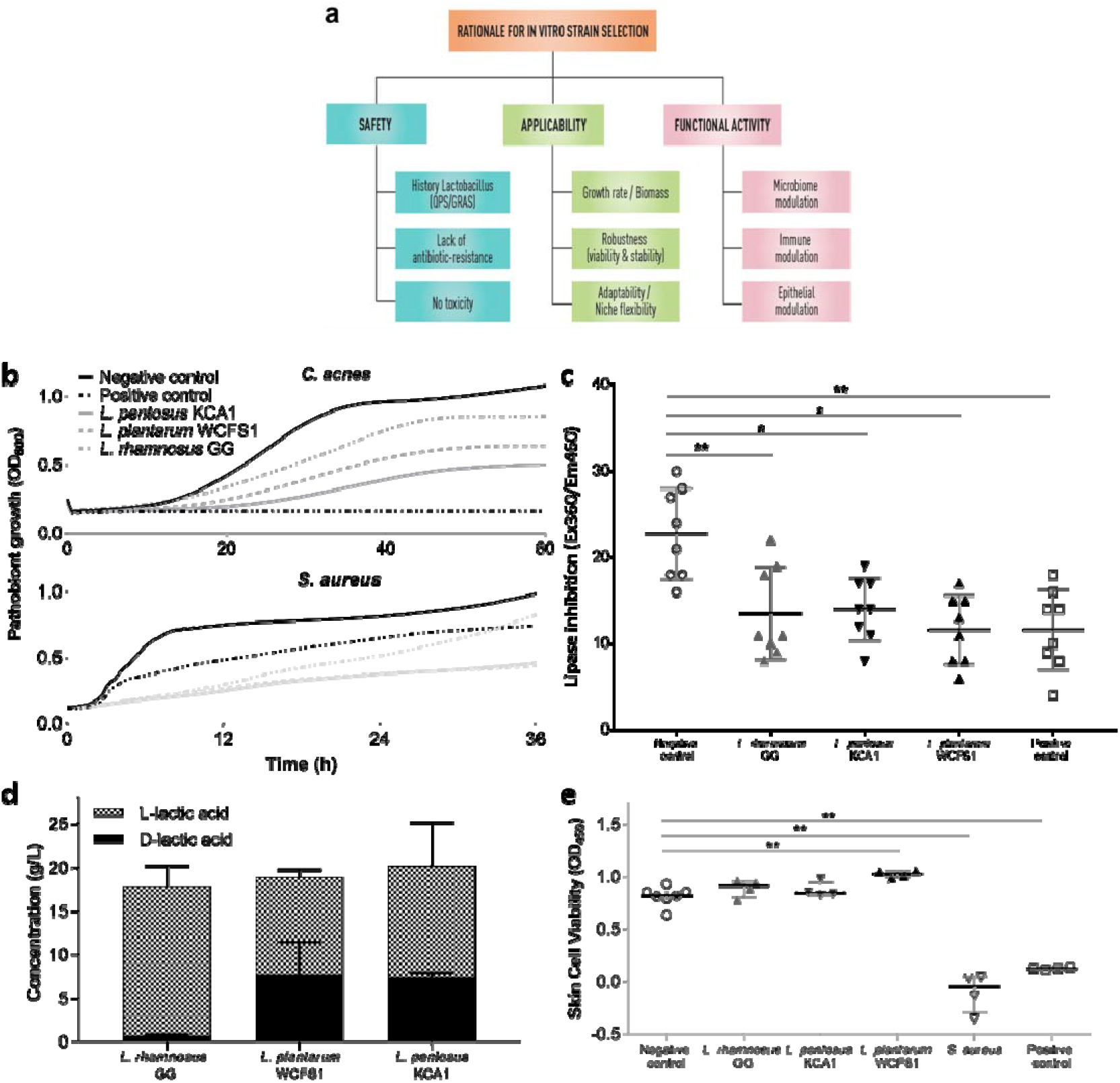
*In vitro* selection of *Lactobacillus* strains for targeted application against acne vulgaris. (a) Schematic overview of the rationale for the selection. Each criterion needs to be taken into account upon selection. Laboratory or genomic prediction tests exist for each criterion. More information can be found in the main text, (b) Antimicrobial activity of the spent-culture supernatant of the selected *Lactobacillus* strains against the two pathobionts tested, *C. acnes* and *S. aureus*, and compared to the positive control (10 mg/mL Clindamycin, a common antibiotic used in acne). MRS at pH4, which is comparable to the pH of the spent supernatant of lactobacilli, was used as a negative control, (c) Inhibition of lipase activity of *C. acnes* by the spent-culture supernatant of the selected *Lactobacillus* strains, and compared to the positive control (10 mg/mL Clindamycin) and the negative control (MRS), (d) Concentration of L-lactic acid and D-lactic acid as key antimicrobial and skin-modulating molecules produced by the selected lactobacilli after overnight incubation in MRS broth, (e) Skin cell viability results of NHEK cells after addition of the selected lactobacilli compared to the negative control, keratinocyte growth medium 2, and positive controls, *S. aureus* and Triton-X, measured at 450 nm using an XTT assay. Statistical analysis were performed using a Mann-Whitney test where * = p<0.05 and ** = p<0.01.

### Viable *Lactobacillus* formulation in O/W cream

We then aimed to design a topical formulation suitable for the application of live bacteria in a sufficient dose on the skin. The selected bacteria were freeze-dried for stability reasons^38^ and embedded in the core of 2-compartment microcapsules (**Figure 3a)**. Various processing conditions were optimized as described in the Methods section and schematized in **Figure 3a**, resulting in capsules of 1500 – 2000 μm diameter with a core of suspended freeze-dried bacteria that can be released upon applying mechanical pressure, such as rubbing on the skin (**Figure 3b)**. Ingredients were selected so that they did not significantly impact on the growth capacity of the skin commensals and pathobionts (tested for *S. epidermis, S. aureus*, and *L. crispatus*) (**Extended Data Figure 2 c-d)**. This formulation and encapsulation approach significantly improved the viability for storage at 4°C and even at 25°C, compared to non-encapsulated freeze-dried bacteria when suspended in a carrier oil-in-water (O/W) cream **(Figure 3c)** and this for up to 6 months (**Figure 3d**).

**Figure 3.**
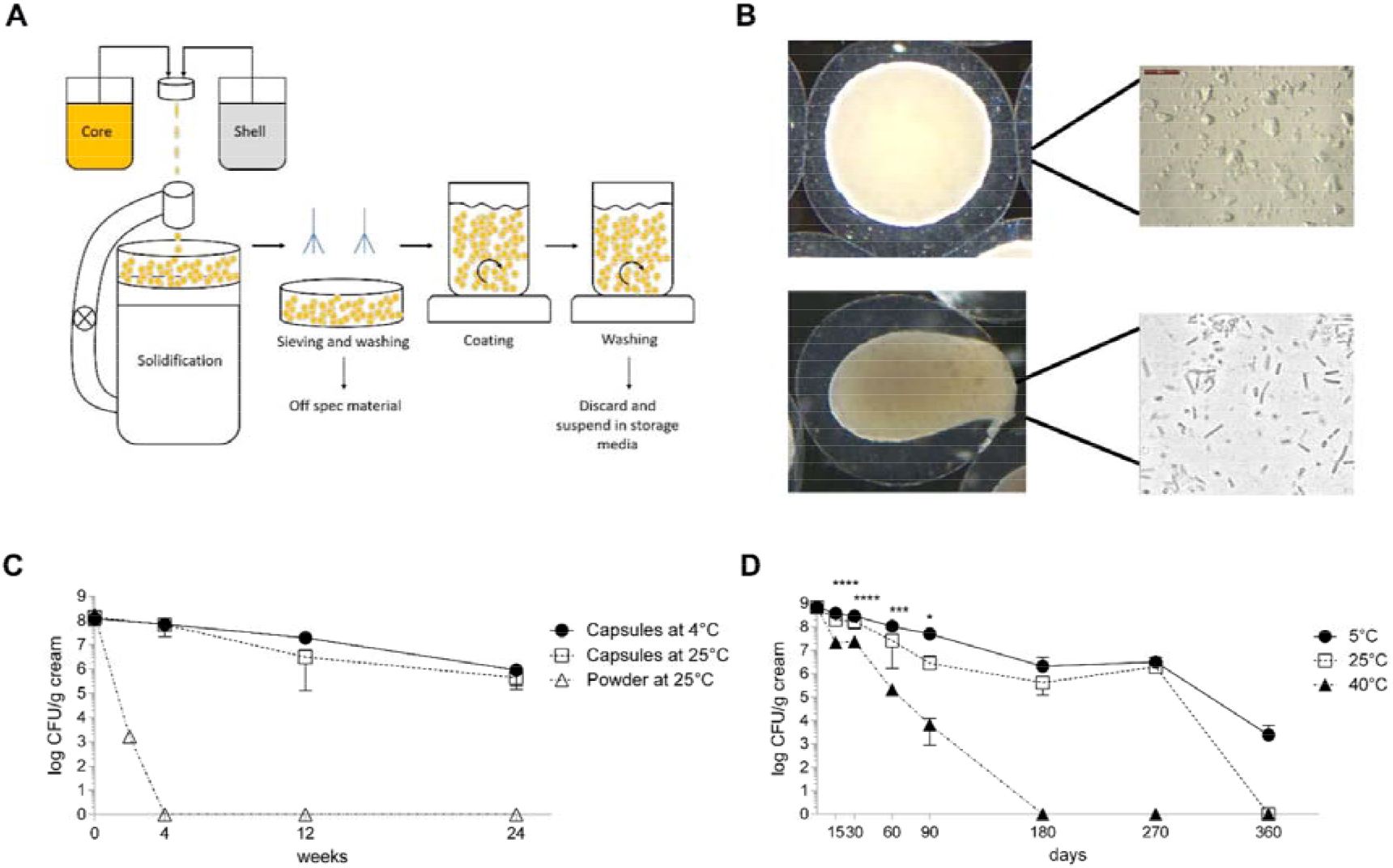
Formulating live lactobacilli in a topical cream. (a) Schematic representation of the micro-encapsulating process with the bacteria in the core suspension and an outer shell made by the shell solution, (b) Resulting micro-capsules with a core of freeze-dried bacteria suspended in oil compared to microcapsules in which a force is applied just before application on the skin, hereby releasing the bacteria and activating them through water-uptake. (c) Survival of the bacteria in the microcapsules after different days compared to nonencapsulated freeze dried bacterial powder in an O/W cream, (d) Survival of the encapsulated bacteria in o/w cream tested according to the International Council for Harmonisation of Technical Requirements for Pharmaceuticals for Human Use Q1A(R2). Statistical analysis was performed using a Two-way ANOVA where * = p<0.05, *** = p<0.001 and **** = p<0.000l.

Subsequently, the skin irritation potential was checked for 20 volunteers with skin patch tests according to Basketter *et al*.^39^. No erythema, dryness or edema was observed in any of the volunteers studied (skin irritation index: 0.00) (**Extended Data Table 2)**. For comparison, adapalene products, which are naphthoic acid derivatives with retinoid activity and documented efficacy in the treatment of mild-to-moderate acne vulgaris, have a mean cumulative irritation index between 0.25–1^40^. Also the widely used combined clindamycin–benzoylperoxide treatment for moderate acne has been reported to frequently induce dry skin, flaky/peeling skin, irritated skin, itchy skin and redness in acne patients^41^.

### *Lactobacillus* skin microbiome modulation

Subsequently, we applied the topical cream twice daily in an open-label ‘ proof-of-concept’ trial to ten volunteers for eight weeks twice daily at 10^8^ CFU per application (± 1 gram/application) (**Figure 4a)**. Patients with mild-to-moderate acne symptoms that were not using antibiotics or another acne treatment were included by the responsible dermatologist (**Extended Data Table 3)**. The impact of the *Lactobacillus* cream on their facial skin microbiome was monitored by 16S amplicon sequencing at four different time points, over a period of 10 weeks (**Figure 4a)**. In this way, the skin baseline microbiome before, during and after the treatment was compared. The skin acne microbiome of these patients at the time of inclusion was especially characterized by an increased relative abundance of *Staphylococcus* taxa (p= 0.0058, Wilcoxon rank sum test) when compared to the healthy controls (**Figure 4b)** (**Extended Data Figure 4** for 3 specific *Staphylococcus* ASVs). No significant difference in relative abundance of *Lactobacillus* taxa was observed between our patients and the reference samples at time of inclusion (**Extended Data Figure 3a-b)**. However, we did observe a significantly reduced relative abundance of *Streptococcus salivarius*, a taxon also belonging to the lactic acid bacteria with lactic acid production as core function (**Extended Data Figure 4)**. After application of the cream with the lactic-acid producing lactobacilli, the facial skin samples of our acne patients at visit 2 and visit 3 clearly clustered separately on a PCoA plot (**Figure 4c)**. Interestingly, in 7 of 10 patients at visit 2 (4 weeks) and 8/10 patients at visit 3 (8 weeks), *Lactobacillus* ASVs were found in relative high abundances (between 20.9% and 92.8%), while in three patients at visit 2 and two patients at visit 3, their relative abundance was below 5% (between 0.015 and 1.1 %) (**Figure 4d and Extended Data Figure 3)**. ASV analysis via EZ taxon^42^ and comparison with the whole genome sequences^22–24^ for rRNA copy variants confirmed that the detected ASVs matched the applied lactobacilli. Interestingly, the three probiotic strains appeared to persist on the skin in similar numbers (**Figure 4d)**. To substantiate that the lactobacilli detected on the skin were still viable, samples were also plated on *Lactobacillus*-selective MRS agar. Most samples at visit 2 (7/9) and visit 3 (6/7) were culture-positive, indicating that – at least some of- the lactobacilli applied were metabolically active on the skin (**Figure 4d)**. At visit 4 (two weeks after the stop of the treatment), most *Lactobacillus* ASVs had disappeared and also growth in MRS medium was markedly reduced, further substantiating that the lactobacilli detected originated from the applied topical cream. We then explored whether the presence of lactobacilli during treatment had impacted on the pathobionts of acne (*C. acnes* and *Staphylococcus* taxa). The relative abundance of both pathobiont taxa dropped indeed at visit 2 and 3 and increased again at visit 4 (p<0.05 for visit 3 versus visit 1 – Wilcoxon test for *Staphylococcus*) (**Figure 4b)**.

**Figure 4.**
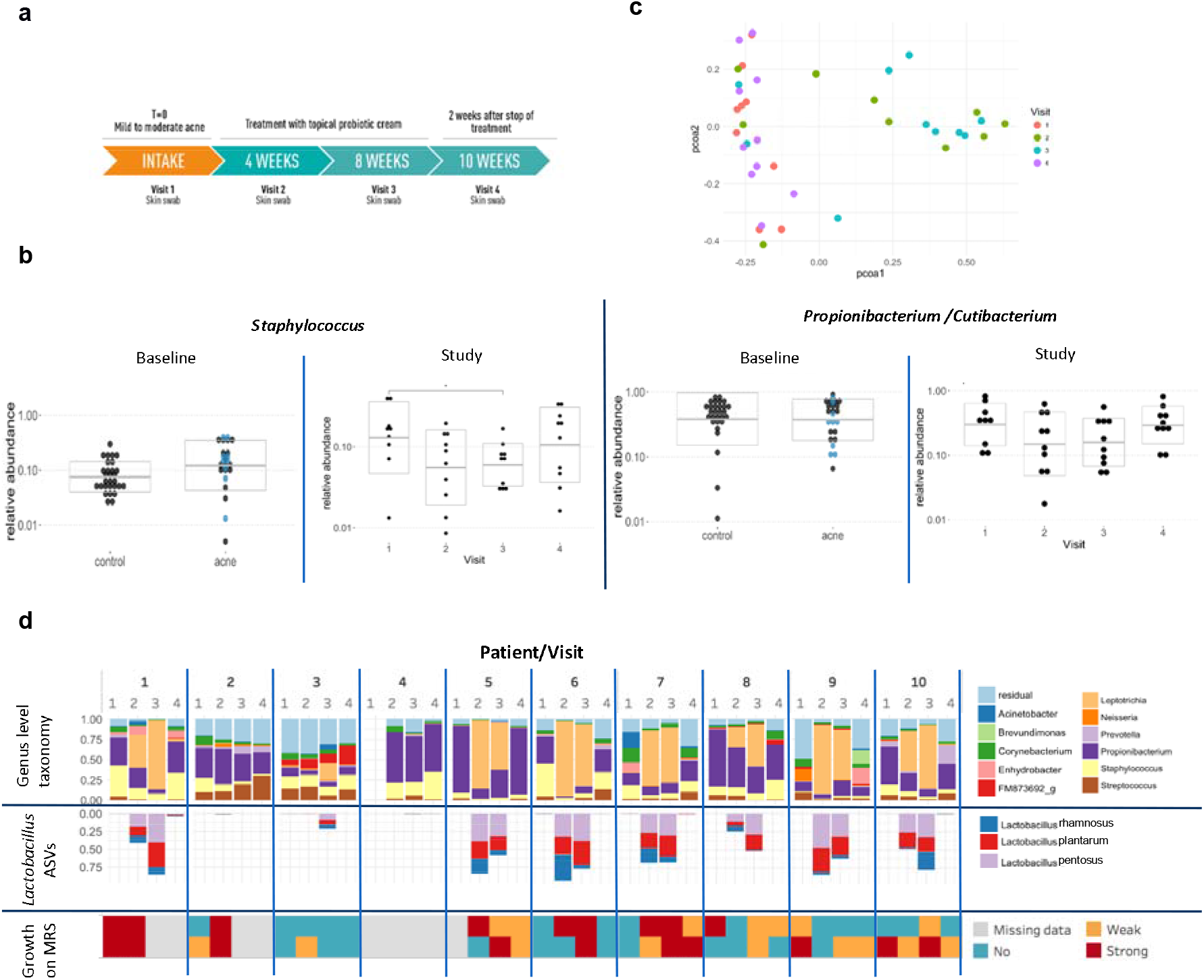
Impact of the *Lactobacillus* cream on the skin microbiome. (a) Schematic overview of the POC intervention study with the O/W cream containing the selected and formulated lactobacilli, the visits at which a skin swab was taken and dermatological symptom analysis was performed by the dermatologist. The cream was applied twice daily for 8 weeks with a minimal dose of 10^6^ CFU/application. (b) Relative abundance of *Staphylococcus* and *Propionibacterium/Cutibacterium* respectively at baseline and over the four visits of the study (right). For the baseline, skin samples of the 30 healthy volunteers without acne symptoms (cfr. Figure 1) and 27 patients with mild-to-moderate acne symptoms were compared. Of these 27 acne patients, 10 patients (indicated with blue dots) were included in the *Lactobacillus* intervention Study (Study) shown at the right side of each panel. For the study visits, p<0.05 for visit 3 versus visit 1 based on Wilcoxon ranks test is indicated with a star, (c) PCOA plot distributing samples according to beta-diversity (Bray-Curtis distance). Similar samples are located closely to each other, and colored by visit, (d) Microbial communities during the study period with the genus-level taxonomy indicated (top), relative abundance of the three *Lactobacillus* ASVs resulting from the cream (middle) and observed growth on MRS medium (top row on agar, bottom row growth in MRS broth) after addition of the skin samples (bottom). Other *Lactobacillus* ASVs were not observed at a higher relative abundance than 1%. Samples were ordered by participant and by visit.

### *Lactobacillus* improvement of acne symptoms

Subsequently, the acne symptoms were clinically scored as the presence of inflammatory lesions and comedones. This analysis showed an overall improvement of the acne symptoms in all patients treated with the *Lactobacillus* cream, as reflected by a significant reduction in inflammatory lesions at visit 2 and 3 compared to visit 1, and a significant reduction in comedone counts at visit 2 (**Figure 5a)**. A significant association between comedonal counts and both *Staphylococcus* and *Propionibacterium/Cutibacterium* was also found (**Figure 5b)**, but not for the inflammatory lesions (**Extended Data Figure 5)**. Of note, when the treatment stopped, the acne scores increased again, indicating that the applied lactobacilli and associated microbiome - staphylococcal modulation – did not persist, in agreement with the fact that the exogenously applied lactobacilli could not permanently colonize (**Figure 4 d)**. On the other hand, this increase in acne scores when the *Lactobacillus* application stopped, further suggests a possible causal association between the applied lactobacilli and the acne symptom reduction.

**Figure 5.**
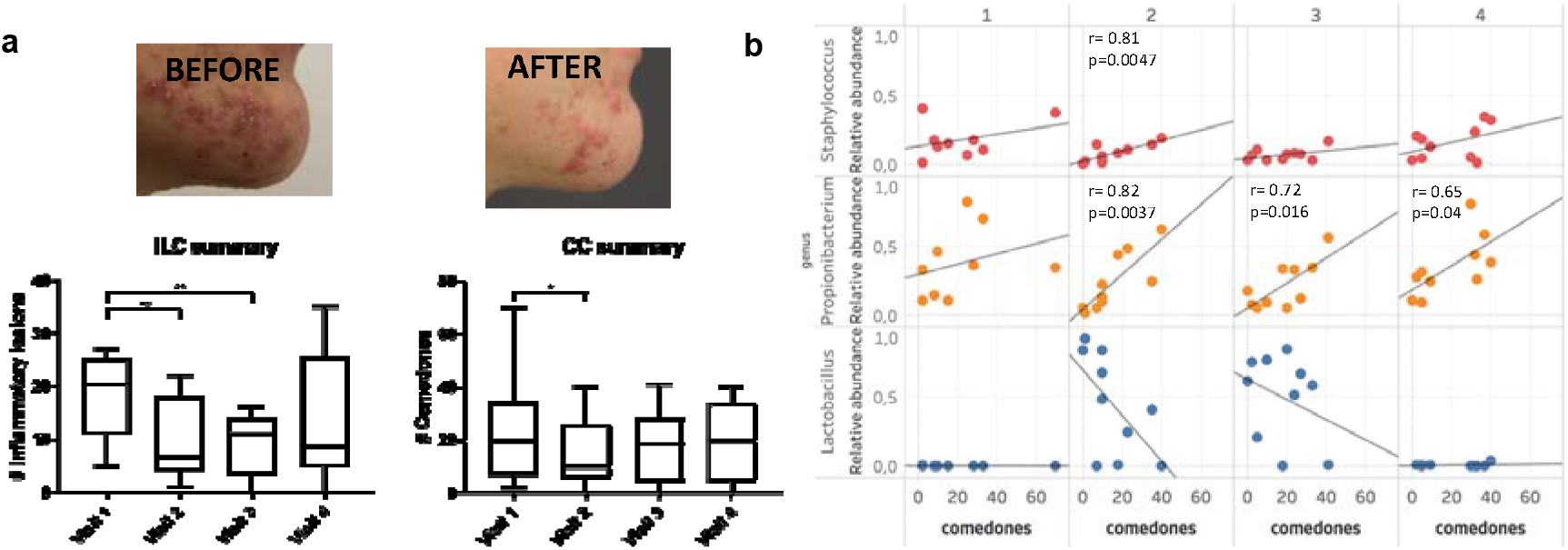
Effect of the *Lactobacillus* cream on acne symptoms and correlation with microbiome data. (a) Evolution of counts of inflammatory lesions (left) and comedones (right) over the course of the study, grouped by visit. All 10 patients included in the pilot study showed a clinical improvement after the application of the cream as exemplified with a picture of the acne spot area of one patients at visit 1 versus visit 2. Statistical analysis were performed using a Wilcoxon matched-pairs signed rank test where * = p<0.05 and ** = p<0.01 (b) Correlation of relative abundances of *Staphylococcus* (top), *Propionibacterium/Cutibacterium* (middle) and *Lactobacillus* (bottom) to comedonal counts, per visit. Pearson correlation coefficient and p-values are indicated where p<0.05.

## Conclusion

Acne vulgaris is a common reason for long-term antibiotic use, with dermatologists prescribing antibiotics more commonly than any other physician group^7^. Here, we applied a multiphasic and multidisciplinary approach to substantiate that *Lactobacillus* strains have potential as skin probiotics to target acne. First, we provided detailed information that lactobacilli (and other lactic acid producing taxa) are unneglectable endogenous members of the human skin microbiota, with relative abundances in between those of human vaginal^43^ and stool^10^ samples. Of interest, phylogenetic placement of the *Lactobacillus* sequences detected in our data (amplicon sequence variants) and the curatedMetagenomicData recently described by Pasolli et al.^13^ showed that the dominant *Lactobacillus* taxa (*L. crispatus, L. iners, L. gasseri, L. jensenii*) of the vaginal community are also among the most prevalent *Lactobacillus* taxa for the skin. Previous studies have briefly acknowledged the presence of lactobacilli in the skin microbiota^44,45^, however such detailed analysis of specific *Lactobacillus* taxa in the skin niche had not yet been performed. Yet, we also showed that to apply selected lactic acid bacteria on the skin, other properties such as robustness to (processing) stress conditions and growth capacity are required, in addition to safety and lack of (transferable) antibiotic resistance properties (as rationalized in **Figure 2a)**. Following this rationalized scheme, we did manage to translate our results directly from *in vitro* lab tests with skin cells and pathogens to human volunteers, without the need for animal testing. Spent-culture supernatant of the selected *L. rhamnosus* GG, *L. plantarum* WCFS1 and *L pentosus* KCA1 could inhibit the growth of *C. acnes* and *S. aureus in vitro*, could survive the formulation in capsules in an O/W cream and were found in similar amounts after 1/1/1 application on the facial skin of patients with mild-to-moderate acne symptoms. Twice daily topical application of this cream with the live lactobacilli was able to reduce inflammatory acne lesions and comedone formation in the ten patients included in the open-label pilot study, and was associated with a reduction in *Staphylococcus* relative abundance (as summarized in **Figure 6)**. Our 16S rRNA ASV-based comparison of the acne facial microbiome of 30 healthy volunteers and 27 patients with acne symptoms suggests indeed that *Staphylococcus* taxa are increased in acne patients and that *Staphylococcus* could thus form an interesting acne target to further investigate. ASV level analysis of the sequenced V4 region of the 16S rRNA gene did not allow identification of the *Staphylococcus* taxa up to species/strain level, so that no distinction between *S. epidermidis* and *s. aureus* and specific more virulent strains could yet be made. On the other hand, microbiome comparison of the skin of subjects with a healthy skin and patients with mild-to-moderate acne vulgaris also pointed to other lactic acid producing bacteria such as *Streptococcus salivarius* as being potentially beneficial against acne.

**Figure 6.**
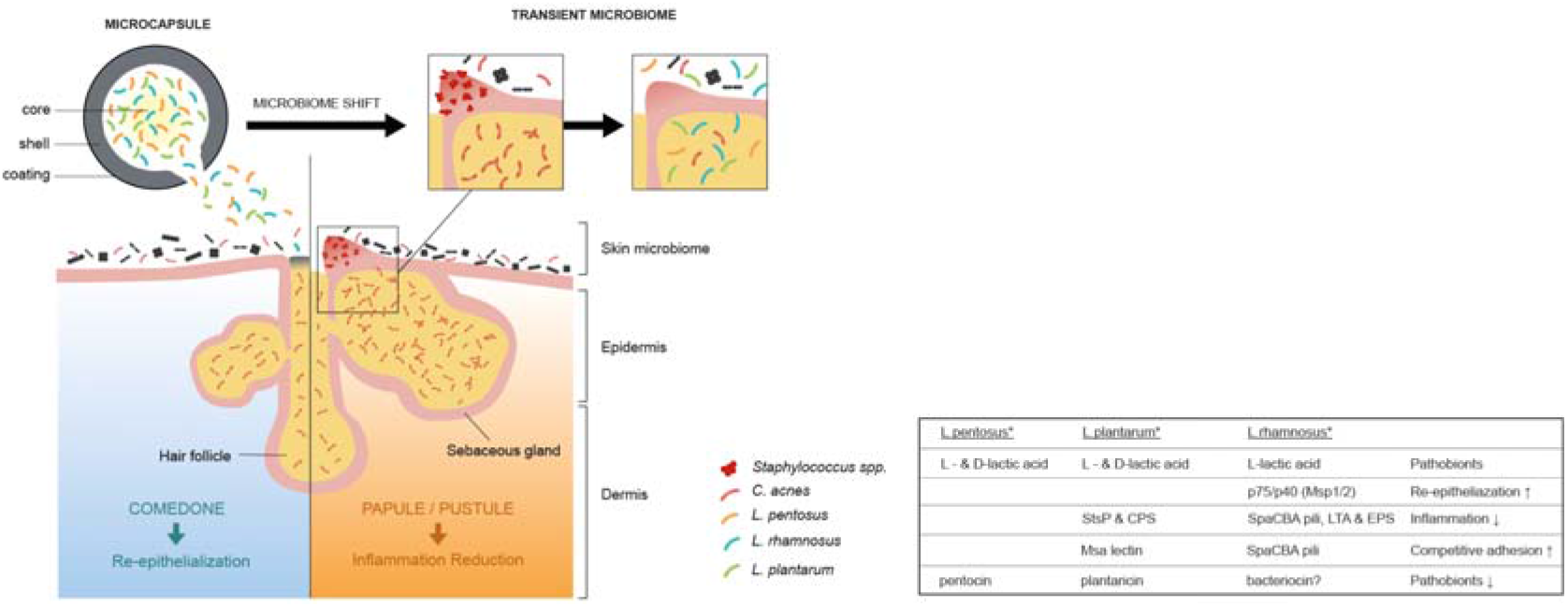
Schematic overview of the main findings of this study on how live lactobacilli formulated in a topical cream modulate skin microbiome and improve acne symptoms. Specific *Lactobacillus* strains were selected and formulated in capsules in an oil (O) in water (W) cream that release the probiotics upon rubbing on the skin. Microbiome analysis (16S amplicon sequencing), as well as counting of comedone and inflammatory lesions substantiated that these lactobacilli could reduce inflammation and comedone formation, as well have transient impact on the skin microbiome, especially by decreasing the relative abundance of staphylococci and *C. acnes* as acne pathobionts. The postulated mode of action (also indicated with * in table) includes their antimicrobial activity against pathobionts by lactic acid (this study), competitive exclusion^47,32^ and possibly bacteriocins^48,24^, their capacity to reduce inflammation, e.g. by the serine-threonine rich protein StsP of *L. plantarum* WCFS1^49^ or SpaCBA pili of *L. rhamnosus* GG^50,51^, and their capacity to promote re-epithelialization^33,34^, by e.g. the secreted proteins Mspl (p75)/Msp2 (p40) for *L. rhamnosus* GG^29,52^. Yet, the involvement of these probiotic effector molecules remains to be further substantiated in follow-up work.

Our findings are consistent with the growing body of evidence that lactic acid bacteria such as lactobacilli can be applied at multiple human body sites to target the microbiome, epithelial barrier function and immune system in various conditions^35^. In this study, we now add support for the skin as topical therapeutic area. Evidently, compliance of this probiotic therapy by the patients will be a key aspect to monitor – and possibly improve-in the future. The promising results from our proof-of-concept study with live lactic acid-producing microbes should now also be confirmed in larger-scale and longitudinal studies, in addition to more molecular studies towards underlying antimicrobial and anti-inflammatory mechanisms and probiotic effector molecules (**Figure 6)**. Together, these studies will contribute to a new era of skin therapeutics based on microbiome modulation, as well as more fundamental and mechanistic insights on the keystone core functions of lactic acid bacteria for skin health.

## Methods

### Bacterial growth

*Lactobacillus* strains were grown at 37°C in de Man, Rogosa and Sharpe (MRS) medium (BD Difco, Erembodegem, Belgium). *Propionibacterium acnes* ATCC6919 was inoculated in reinforced clostridial broth (LabM Limited, Heywood, UK), supplemented with 0.2% Tween20 and cultured microaerobically (5% CO_2_) at 37°C. *Staphylococcus aureus* ATCC29213 was grown in Mueller-Hinton broth at 37°C. Solid media contained 1.5% (w/v) agar. Time-course experiments were also performed analysing the antimicrobial activity of spent culture supernatant (SCS) of the selected *Lactobacillus* strains against C. *acnes* and *S. aureus* ATCC29213 (cfr.^53^). Additionally, the impact of this SCS (10%) on the lipase activity of *C. acnes* was determined as previously described ^5^.

### Human skin cell culture

Normal human epidermal keratinocytes (NHEK) cells from juvenile foreskin from pooled donors were purchased from Promocell (Heidelberg, Germany) and cultured according to manufacturer’s recommendations in Keratinocyt Growth medium 2 (Promocell, Heidelberg, Germany). Cytotoxicity of probiotic strains was assessed using the 2,3-Bis(2-methoxy-4-nitro-5-sulfophenyl)-2H-tetrazolium-5-carboxanilide (XTT, Sigma-Aldrich) cell viability assay. NHEK cells were seeded at a density of 5000 cells/well in 96-well plates and cultured until confluent. Overnight cultures of probiotic strains or *S. aureus* were added to the wells with or without NHEK cells at 10^6^ CFU/well and incubated for 2 h at 5% CO_2_ and 37°C. Triton X-100 (0.5%) was used as a positive control.

### Collection of skin samples and total microbial DNA extractions

Skin samples were collected by brushing the cheek (control group) or the affected area on the face (patients) with a FloqSwab (Copan) over an area of ± 10 cm^2^ or around the lesions. Swabs were then transferred to a falcon containing 800μl Bead solution of QIAamp PowerFecal DNA kit (Qiagen). Samples were stored at 4°C until further processing (maximally 14 days). Before DNA extraction, samples were vortexed for 1 minute, after which the Bead solution was transferred to the bead tube. Subsequent steps of the DNA extraction were executed according to manufacturer’s instructions.

### Illumina MiSeq *16S rDNA* gene amplicon sequencing

The primers used for Illumina MiSeq sequencing were based on the previously described 27F-338R or 515F-806R primers^54^ and altered for dual-index paired-end sequencing, as described earlier^55^ (**Extended Data Table 4**). Separate runs were carried out for V1V2 and V4 *rRNA* gene variable regions. Quality control and processing of reads was performed using the R package DADA2, version 1.6.0 ^56^. Denoised reads (amplicon sequence variants or ASVs) were merged and read pairs with one or more conflicting bases between the forward and reverse read were removed. Chimeric sequences were removed using the function “removeBimeraDenovo”. Finally, ASVs were classified from the kingdom to the genus level using the EzBioCloud 16S database^42^. A species annotation was added to each ASV by listing the species of all 16S sequences in the database that showed an exact match to the ASV sequence. Contaminants were identified using the approach of Jervis-Bardy *et al*^57^. ASVs with a strong negative correlation between relative abundances and total sample read counts were considered contamination. For each ASV, this correlation was calculated and tested for significance. ASVs with a p-value less than 0.0001 were removed. Samples were filtered by removing those with less than 1000 reads left after all read and ASV filtering steps.

### Biostatistical and bioinformatics analysis

Processing of the ASV table, ASV annotations (e.g. classification) and sample annotations (metadata) were performed using the in-house R package “tidyamplicons”, publicly available at github.com/SWittouck/tidyamplicons. For the analyses at the genus level, ASV read counts were aggregated at the genus level or, if unavailable, at the most specific level at which taxonomic annotation was available.

### Analysis of public datasets

Processed OTU-table and sample metadata from the Human Microbiome Project (HMPv35)^46^ and the shotgun metagenomic datasets were retrieved using the MicrobeDS R package and curatedMetagenomics R package^16^ respectively. All data was loaded, processed and visualized in the R-environment using Phyloseq. All scripts are available at https://github.com/LebeerLab/skin_acne_study.

### qPCR for estimation of absolute bacterial concentrations

qPCR was performed in duplicate on a 20-fold dilution (to avoid interference of PCR inhibitors) of total DNA isolated from the samples, using the StepOnePlus real time qPCR system (Applied Biosystems^®^, Foster City, California, USA), SYBR^®^ Green chemistry (PowerUp^™^ SYBR^®^ Green Master Mix, Applied Biosystems^®^, Foster City, California, USA) and primers as indicated in **Extended Data Table 4**. Standard curves were used to estimate bacterial concentrations in the samples and derived from serially diluted DNA from an overnight culture of *Lactobacillus crispatus* LMG12005 isolated similarly as the samples. Bacterial concentration was determined by plating.

### Formulation of lactobacilli in microcapsules and O/W cream

A single colony of the three selected probiotic strains was grown until stationary phase and lyophilized. The lyophilized bacterial powder was grinded and milled (Frewitt, Switzerland) to obtain a fine powder (± 10^11^ CFU/gram) and subsequently encapsulated via a core-shell encapsulation approach. Briefly, the strains were mixed in equal amounts and homogeneously suspended to obtain a stable oil-based feed core suspension. The shell feed solution contained a hydrocolloid alginate polymer as gelling agent. Both liquid feeds were pumped to a concentric nozzle, to obtain a concentric fluid flow. The laminar liquid flow was broken up by a vibrational unit to obtain spherical -droplets that were solidified upon falling in a calcium-based solidification solution, forming the capsules. The collected capsules (10^9^ - 10^10^ CFU/gram) were washed and suspended in an oil-in-water cream containing the following ingredients: Aqua, Glycerin, Polyglyceryl-3 Rice Branate, Propylheptyl Caprylate, Cetyl Alcohol, Nylon 6/12, Caprylic/Capric Triglyceride, Squalane, PEG-400, Cyclopentasiloxane, Cetearyl Alcohol, Prunus Amygdalus Dulcis Oil, Allantoin, Limnanthes Alba Seed Oil, Tocopheryl Acetate, Xanthan Gum, Cyclohexasiloxane, Helianthus Annuus Seed Oil, Algin, p-Anisic Acid, Silica Dimethyl Silylate, Disodium EDTA, Sodium Hydroxide, Hydrochloric Acid. The ingredients of this cream, mainly the emulsifiers and preservatives, were selected to be compatible with the micro-capsules and bacteria, both during storage and upon release of the probiotics. Hereto, the impact of the topical cream without the capsules on the growth of four skin reference bacteria (S. *aureus, S. epidermidis, L. crispatus* and *C.acnes*) was evaluated at a concentration of 1, 10 and 100 mg/ml, by a time-course analysis of OD600 measurements as described above. Mechanical force (rubbing on the skin) was confirmed to break the capsules, releasing the inner core material containing the suspended probiotics. Skin irritation tests with the O/W cream containing the freeze –dried lactobacilli were performed as described previously^39^.

### Proof-of-concept human in patients with acne vulgaris

A proof-of-concept clinical trial was performed on patients with mild-to-moderate acne vulgaris included after careful assessment of the responsible dermatologist by counting of comedones and inflammatory lesions (**Extended Data Table 3**). Patients were men between 12–25 years. Exclusion criteria were use of oral antibiotics within 4 weeks prior to start of the study and use of systemic retinoids within 6 months prior to start of study. Subjects provided written informed consent before the study began. Patients were asked to apply the topical probiotic cream (containing 10^8^ CFU of each *Lactobacillus* strain per application of 1 g of the topical cream). Patients were asked to apply the cream twice daily for 56 days (8 weeks). The patients were seen by a dermatologist at start (before the therapy) (visit 1), week 4 (visit 2), week 8 (visit 3) and week 10 (visit 4). A skin swab was taken at each visit, total DNA was extracted and amplified for 16S amplicon sequencing as described above. Moreover, a clinical scoring was performed and a photograph taken at each visit.

## Clinical trial registration

The protocol of this study was in accordance with the Declaration of Helsinki, and was approved by the ethics committee of the University Hospital of Antwerp (Belgium) before initiation of the study. The study was given the approval number B300201628507 (Belgian registration) and registered online at clinicaltrials.gov with unique identifier NCT03469076.

## Data availability

Sequencing data are available at the European Nucleotide Archive with the accession number PRJEB27311.

**Extended Data** is available in the online version of this manuscript.

## Acknowledgements

We kindly acknowledge Dr. K. Anukam for providing us *Lactobacillus pentosus* KCA1. We thank Wannes Van Beeck, Dieter Vandenheuvel, Camille Allonsius, Leen Van Ham and all other members of the ENdEMIC group for their assistance and/or fruitful discussions. We thank S. Van Elslander for the help in the design of figures 2A, 4A and 6. We also greatly thank Dr. E. Pasolli for the assistance with the CuratedMetagenomics Package. This work was funded by grants from the Flanders Agency for Innovation & Entrepreneurship (www.vlaio.be/en), including an IWT-SBO project (IWT/50052) for the fundamental part of the research and a VLAIO R&D project with Yun Probiotherapy NV (formerly known as Axca Bvba) as SME partner for the applied part of the project. The qPCR analyses were performed with a StepOne Plus (Applied Biosystems) machine funded by the Fund for Scientific Research Flanders (1520114N). Sander Wuyts and Ilke De Boeck hold a personal PhD grant (IWT-SB 141198 and FWO 1S17916N respectively).

## Author contributions

S.L., I.C., J.L. designed the study; J.L. clinically evaluated the patients and collected patient samples; E.O. prepared the clinical and control samples for MiSeq sequencing with the help of I.T. and I.D.B., Sa.Wu. did the shotgun metagenome analysis, Sa.Wu. St.Wi., and E.O. analyzed sequence data. E.O., M.V.B., C.A. and I.S. did part of the microbiological and cell culture lab experiments. I.C., T.H., F.K. formulated the lactobacilli in the topical cream. S.L. drafted the manuscript and all authors approved the manuscript.

## Author information

The authors declare the following competing interests. I.C. and T.H. were employed at UAntwerp at the time of the study, but are currently working at the R&D department of Yun NV, a start-up company resulting from this research (www.yun.be). Based on the data presented here, YUN NV has selected and formulated three *Lactobacillus* strains, *L. pentosus* YUN-V1.0, *L. plantarum* YUN-V2.0 and *L. rhamnosus* YUN-S1.0 in their commercial ACN product. Part of the results presented in this manuscript are included in patent applications PCT/EP2017/066176 and PCT/EP2017/065006.

## References

1. Grice, E. A. et al. Topographical and temporal diversity of the human skin microbiome. Science (80-.). 324,1190–1192 (2009).

2. Byrd, A. L., Belkaid, Y. & Segre, J. A. The human skin microbiome. Nature Reviews Microbiology 16,143–155 (2018).

3. Hill, C. et al. Expert consensus document: The International Scientific Association for Probiotics and Prebiotics consensus statement on the scope and appropriate use of the term probiotic. Nat. Rev. Gastroenterol. Hepatol. 11, 506–514 (2014).

4. Scholz, C. F. P. & Kilian, M. The natural history of cutaneous propionibacteria, and reclassification of selected species within the genus propionibacterium to the proposed novel genera acidipropionibacterium gen. Nov., cutibacterium gen. nov. and pseudopropionibacterium gen. nov. Int. J. Syst. Evol. Microbiol. 66, 4422–4432 (2016).

5. Higaki, S. Lipase inhibitors for the treatment of acne. Journal of Molecular Catalysis B: Enzymatic 22, 377–384 (2003).

6. Dreno, B. What is new in the pathophysiology of acne, an overview. Journal of the European Academy of Dermatology and Venereology (2017). doi:10.1111/jdv.14374

7. Del rosso, J. Q. et al. Status report from the scientific panel on antibiotic use in dermatology of the American acne and rosacea society part 1: Antibiotic prescribing patterns, sources of antibiotic exposure, antibiotic consumption and emergence of antibiotic resistance, impac. J. Clin. Aesthet. Dermatol. (2016).

8. Williams, H. C., Dellavalle, R. P. & Garner, S. Acne vulgaris, in The Lancet 379, 361–372 (2012).

9. Bernardeau, M., Guguen, M. & Vernoux, J. P. Beneficial lactobacilli in food and feed: Long-term use, biodiversity and proposals for specific and realistic safety assessments. FEMS Microbiology Reviews (2006). doi:10.1111/j.1574-6976.2006.00020.x

10. Heeney, D. D., Gareau, M. G. & Marco, M. L. Intestinal Lactobacillus in health and disease, a driver or just along for the ride? Current Opinion in Biotechnology 49,140–147 (2018).

11. Petrova, M. I., Lievens, E., Malik, S., Imholz, N. & Lebeer, S. Lactobacillus species as biomarkers and agents that can promote various aspects of vaginal health. Frontiers in Physiology 6, (2015).

12. Martensson, A. etal. Effects of a honeybee lactic acid bacterial microbiome on human nasal symptoms, commensals, and biomarkers. Int. Forum Allergy Rhinol. 6, 956–963 (2016).

13. Pasolli, E. et al. Accessible, curated metagenomic data through ExperimentHub. Nature Methods 14,1023–1024 (2017).

14. Chng, K. R. et al. Whole metagenome profiling reveals skin microbiome-dependent susceptibility to atopic dermatitis flare. Nat. Microbiol. 1, (2016).

15. Oh, J. et al. Biogeography and individuality shape function in the human skin metagenome. Nature 514, 59–64 (2014).

16. Tett, A. et al. Unexplored diversity and strain-level structure of the skin microbiome associated with psoriasis, npj Biofilms Microbiomes 3,14 (2017).

17. Huttenhower, C. & Human Microbiome Project Consortium. Structure, function and diversity of the healthy human microbiome. Nature (2012). doi:10.1038/naturell234

18. Wuyts, S. et al. Carrot juice fermentations as man-made microbial ecosystems dominated by lactic acid bacteria. Appl. Environ. Microbiol. (2018). doi:10.1128/AEM.00134-18

19. Duar, R. M. et al. Lifestyles in transition: evolution and natural history of the genus Lactobacillus. FEMS Microbiol. Rev. 41, S27–S48 (2017).

20. Chu, D. M. et al. Maturation of the infant microbiome community structure and function across multiple body sites and in relation to mode of delivery. Nat. Med. 23, 314–326 (2017).

21. Banerjee, S., Schlaeppi, K. & van der Heijden, M. G. A. Keystone taxa as drivers of microbiome structure and functioning. Nature Reviews Microbiology (2018). doi:10.1038/s41579-018-0024-1

22. Kankainen, M. et al. Comparative genomic analysis of Lactobacillus rhamnosus GG reveals pili containing a human-mucus binding protein. Proc. Natl. Acad. Sci. U. S. A. 106, (2009).

23. Kleerebezem, M. et al. Complete genome sequence of Lactobacillus plantarum WCFS1. Proc. Natl. Acad. Sci. 100,1990–1995 (2003).

24. Anukam, K. C. et al. Genome Sequence of Lactobacillus pentosus KCA1: Vaginal Isolate from a Healthy Premenopausal Woman. PLoS One 8, e59239 (2013).

25. Segers, M. E. & Lebeer, S. Towards a better understanding of Lactobacillus rhamnosus GG – host interactions. Microb. Cell Fact. 13, S7 (2014).

26. van den Nieuwboer, M., van Hemert, S., Claassen, E. & de Vos, W. M. Lactobacillus plantarum WCFS1 and its host interaction: a dozen years after the genome. Microbial Biotechnology 9, 452–465 (2016).

27. Tapiovaara, L. et al. Absence of adverse events in healthy individuals using probiotics – analysis of six randomised studies by one study group. Benef. Microbes 7,161–169 (2016).

28. van Baarlen, P. et al. Differential NF-kB pathways induction by Lactobacillus plantarum in the duodenum of healthy humans correlating with immune tolerance. Proc. Natl. Acad. Sci. (2009). doi:10.1073/pnas.0809919106

29. van Baarlen, P. et al. Human mucosal in vivo transcriptome responses to three lactobacilli indicate how probiotics may modulate human cellular pathways. Proc. Natl. Acad. Sci. U. S. A. 108 Suppl, 4562–4569 (2011).

30. Skovbjerg, S. et al. Spray bacteriotherapy decreases middle ear fluid in children with secretory otitis media. Arch. Dis. Child. 94, 92–98 (2009).

31. Reid, G. & Bruce, A. W. Selection of Lactobacillus Strains for Urogenital Probiotic Applications. J. Infect. Dis. 183, 77–80 (2001).

32. Mohammedsaeed, W., McBain, A. J., Cruickshank, S. M. & O’Neill, C. A. Lactobacillus rhamnosus GG inhibits the toxic effects of Staphylococcus aureus on epidermal keratinocytes. Appl. Environ. Microbiol. 80, 5773–5781 (2014).

33. Mohammedsaeed, W., Cruickshank, S., McBain, A. J. & O’Neill, C. A. Lactobacillus rhamnosus GG Lysate Increases Re-Epithelialization of Keratinocyte Scratch Assays by Promoting Migration. Sci. Rep. 5, (2015).

34. O’Neill, C. A., Sultana, R. & McBain, A. J. Strain-dependent augmentation of tight-junction barrier function in human primary epidermal keratinocytes by lactobacillus and bifidobacterium lysates. Appl. Environ. Microbiol. 79, 4887–4894 (2013).

35. Sanders, M. E., Benson, A., Lebeer, S., Merenstein, D. J. & Klaenhammer, T. R. Shared mechanisms among probiotic taxa: implications for general probiotic claims. Curr. Opin. Biotechnol. 49, (2018).

36. Tang, S. C. & Yang, J. H. Dual effects of alpha-hydroxy acids on the skin. Molecules 23, (2018).

37. European Food Safety Authority. Guidance on the assessment of bacterial susceptibility to antimicrobials of human and veterinary importance. EFSAJ. (2012).doi: 10.2903/j.efsa.2012.2740.

38. Broeckx, G., Vandenheuvel, D., Claes, I. J. J., Lebeer, S. & Kiekens, F. Drying techniques of probiotic bacteria as an important step towards the development of novel pharmabiotics. Int.J. Pharm. 505, (2016).

39. Basketter, D. A., Whittle, E., Griffiths, H. A. & York, M. The identification and classification of skin irritation hazard by a human patch test. Food Chem. Toxicol. 32, 773–775 (1994).

40. Queille-Roussel, C. et al. Comparison of the cumulative irritation potential of adapalene gel and cream with that of erythromycin/tretinoin solution and gel and erythromycin/isotretinoin gel. Clin. Ther. 23, 205–212 (2001).

41. Feldman, S. R. & Chen, D. M. How Patients Experience and Manage Dryness and Irritation From Acne Treatment. J Drugs Dermatol (2011).

42. Yoon, S. H. et al. Introducing EzBioCloud: A taxonomically united database of 16S rRNA gene sequences and whole-genome assemblies. Int. J. Syst. Evol. Microbiol. (2017). doi: 10.1099/ijsem.0.001755

43. Petrova, M. I., Lievens, E., Malik, S., Imholz, N. & Lebeer, S. Lactobacillus species as biomarkers and agents that can promote various aspects of vaginal health. Front. Physiol. 6, (2015).

44. Zeeuwen, P. L. J. M. et al. Microbiome dynamics of human epidermis following skin barrier disruption. Genome Biol. (2012). doi:10.1186/gb-2012-13-11-r101

45. Li, X., Yuan, C., Xing, L. & Humbert, P. Topographical diversity of common skin microflora and its association with skin environment type: An observational study in Chinese women. Sci. Rep. (2017). doi:10.1038/s41598-017-18181-5

46. Human, T. & Project, M. Structure, function and diversity of the healthy human microbiome. Nature (2012). doi:10.1038/nature11234

47. Tytgat, H. L. P. et al. Lactobacillus rhamnosus GG Outcompetes Enterococcus faecium via Mucus-Binding Pili: Evidence for a Novel and Heterospecific Probiotic Mechanism. Appl. Environ. Microbiol. 82, 5756–5762 (2016).

48. Diep, D. B., Straume, D., Kjos, M., Torres, C. & Nes, I. F. An overview of the mosaic bacteriocin pin loci from Lactobacillus plantarum. Peptides (2009). doi:10.1016/j.peptides.2009.05.014

49. Remus, D. M., Kleerebezem, M. & Bron, P. A. An intimate tête-à-tête - How probiotic lactobacilli communicate with the host, in European Journal of Pharmacology 668, (2011).

50. Lebeer, S. et al. Functional analysis of lactobacillus rhamnosus GG pili in relation to adhesion and immunomodulatory interactions with intestinal epithelial cells. Appl. Environ. Microbiol. 78, (2012).

51. Vargas García, C. E. et al. Piliation of Lactobacillus rhamnosus GG promotes adhesion, phagocytosis, and cytokine modulation in macrophages. Appl. Environ. Microbiol. 81, (2015).

52. Yan, F. etal. A lactobacillus rhamnosus GG-derived soluble protein, p40, stimulates ligand release from intestinal epithelial cells to transactivate epidermal growth factor receptor. J. Biol. Chem. 288, 30742–30751 (2013).

53. van den Broek, M. F. L. Multifactorial inhibition of lactobacilli against the respiratory tract pathogen Moraxella catarrhalis. Benef. Microbes 9, 429–439 (2018).

54. Caporaso, J. G. et al. Global patterns of 16S rRNA diversity at a depth of millions of sequences per sample. Proc Natl Acad Sci USA 108, (2010).

55. Kozich, J. J., Westcott, S. L., Baxter, N. T., Highlander, S. K. & Schloss, P. D. Development of a dual-index sequencing strategy and curation pipeline for analyzing amplicon sequence data on the MiSeq Illumina sequencing platform. Appl. Environ. Microbiol. 79, 5112–20 (2013).

56. Callahan, B. J. et al. DADA2: High-resolution sample inference from Illumina amplicon data. Nat. Methods 13, 581–583 (2016).

57. Jervis-Bardy, J. et al. Deriving accurate microbiota profiles from human samples with low bacterial content through post-sequencing processing of Illumina MiSeq data. Microbiome (2015). doi:10.1186/s40168-015-0083-8

